# Non-enzymatic isothermal strand displacement and amplification (NISDA) does not enable sensitive nucleic acid quantification

**DOI:** 10.1101/2025.05.19.654176

**Authors:** Thijs Van der Snickt, Scott Ailliet, Simone Bassini, Anouk Peymen, Ahmet Colaker, Enrico Cadoni, Alejandro Valverde, Elise Daems, Annemieke Madder, Karolien De Wael, Pieter Mestdagh

**Affiliations:** OncoRNALab, Department of Biomolecular Medicine, Ghent University, 9000 Ghent, Belgium; Antwerp engineering, photoelectrochemistry and sensing (A-PECS), Department of Bioscience Engineering, University of Antwerp, 2020 Antwerp, Belgium; NANOlab Center of Excellence, University of Antwerp, 2020 Antwerp, Belgium; OBCR Group, Department of Organic and Macromolecular Chemistry, Ghent University, 9000 Ghent, Belgium

## Abstract

Enzyme-free isothermal amplification methods offer a promising alternative to enzymatic assays for nucleic acid detection, particularly in low-resource settings. The nonenzymatic isothermal strand displacement and amplification (NISDA) assay was recently introduced as a highly sensitive, enzyme-free detection strategy. Here, we attempted to replicate its reported performance. Despite extensive testing, we failed to replicate the reported sensitivity and could only produce detectable signals at extremely high target concentrations (≥1 × 10^11^ copies/µL). Our results highlight the critical importance of independent validation in the development of nonenzymatic diagnostic assays.

## Introduction

Enzyme-free nucleic acid detection methods are gaining traction as low-cost, robust alternatives to traditional enzymatic assays like RT-qPCR and NGS. These conventional methods, while sensitive, depend on enzymes that are costly, temperature-sensitive, and prone to supply chain disruptions. In contrast, isothermal, nonenzymatic strategies such as hybridization chain reaction (HCR)^1^, catalytic hairpin assembly (CHA)^2^ or entropy-driven catalysis (EDC)^3^ offer a simpler and more cost-effective alternative for nucleic acid detection.

Building on this concept, Mohammadniaei et al. introduced the nonenzymatic isothermal strand displacement and amplification (NISDA) assay. A one-pot, enzyme-free system that combines strand displacement and hybridization-driven signal amplification^4^. NISDA was reported to detect long RNA targets with high sensitivity under isothermal conditions, making it highly appealing for integration into point-of-care or label-free sensing platforms.

Motivated by its potential, we sought to adapt the NISDA method within a bio-electrochemical sensor to improve sensitvity. However, despite following the published protocol and testing multiple variables, we were unable to replicate the reported sensitivity. Here, we present our experimental findings and discuss their broader implications for the development and application of nonenzymatic amplification strategies in molecular diagnostics.

## Materials and methods

TES buffer (50 mM Tris-HCl, pH 7.8, 1 mM EDTA, 50 mM NaCl) was prepared for use in the experiments. M1 and M2 probes were obtained from two providers, Integrated DNA Technologies (IDT) and Pentabase, and reconstituted from a lyophilized stock solution (300 µM in water) to a working solution (100 µM in TES buffer). M1 and M2 probes were folded using three different methods: RT (incubation at 95°C for 5 min. followed by cooling to room temperature), 1s (incubation at 95°C followed by ramped cooling to room temperature at 0.1°C/s and a 1-hour incubation at room temperature), and 4s (incubation at 95°C followed by ramped cooling to room temperature at 0.1°C/s with intermediate stops at 77°C, 59°C, and 41°C for 12 minutes each). The initiator (Pentabase) and template oligo (IDT) were obtained lyophilized and reconstituted from stock solutions (300 µM in water) to a working solution (200 µM in TES buffer). A mixture containing the template and initiator (T/INA) was prepared and incubated at 95°C for 5 minutes followed by cooling to room temperature. Concentrations of M1, M2, template, and template/initiator complex were measured using a Nanodrop 1000 Spectrophotometer. Mixtures of M1 at 22 ng/µL, M2 at 85 ng/µL, and the template/initiator complex at 380 ng/µL were prepared. A master mix with 10 µL M1, 10 µL M2, and 4 µL template/initiator or 4µL water when the assay performance was tested on the template was created. 5µL of diluted synthetic template or target was added to run the assay. Hereafter, relative fluorescence unit (RFU) measurements were performed using a CFX Real-Time PCR instrument (Bio-Rad). Thermal cycler protocol involved a 2-min. incubation at 42°C, followed by a 10-second capture at 42°C for 15 cycles with a lid temperature of 43°C. For the data analysis, no baseline correction was performed. The graphs are normalized against their minimum values. Depending on the experiment, the focus was on the amplification part or the total NISDA assay, involving dilution of the template/target as required.

## Results

To evaluate the performance and reproducibility of both the amplification and displacement steps, a dilution series was prepared for the template (**Fig. 1a**) and N-gene target (**Fig. 1b**), according to the protocol mentioned in the original paper (**<u>Materials and methods</u>**). We observed a difference in endpoint fluorescence between the template and no template controls for template concentrations ≥ 1 x 10^11^ copies/µL (**Fig. 1a**). Template concentrations < 1 x 10^11^ copies/µL did not result in a discernable change in endpoint fluorescence. Similar results were obtained when experiments were performed by another operator in another lab (**Supplementary Fig. 1**) and when using M1 and M2 probes synthesized by a different provider (**Supplementary Fig. 2**). Moreover, when verifying the full NISDA assay workflow with a serial dilution of N-gene target, no discernable difference in endpoint fluorescent was observed for target concentrations below 1 x 10^12^ copies/µL (**Fig. 1b**), irrespective of the M1/M2 probe set (**Supplementary Fig. 2**).

**Fig. 1.**
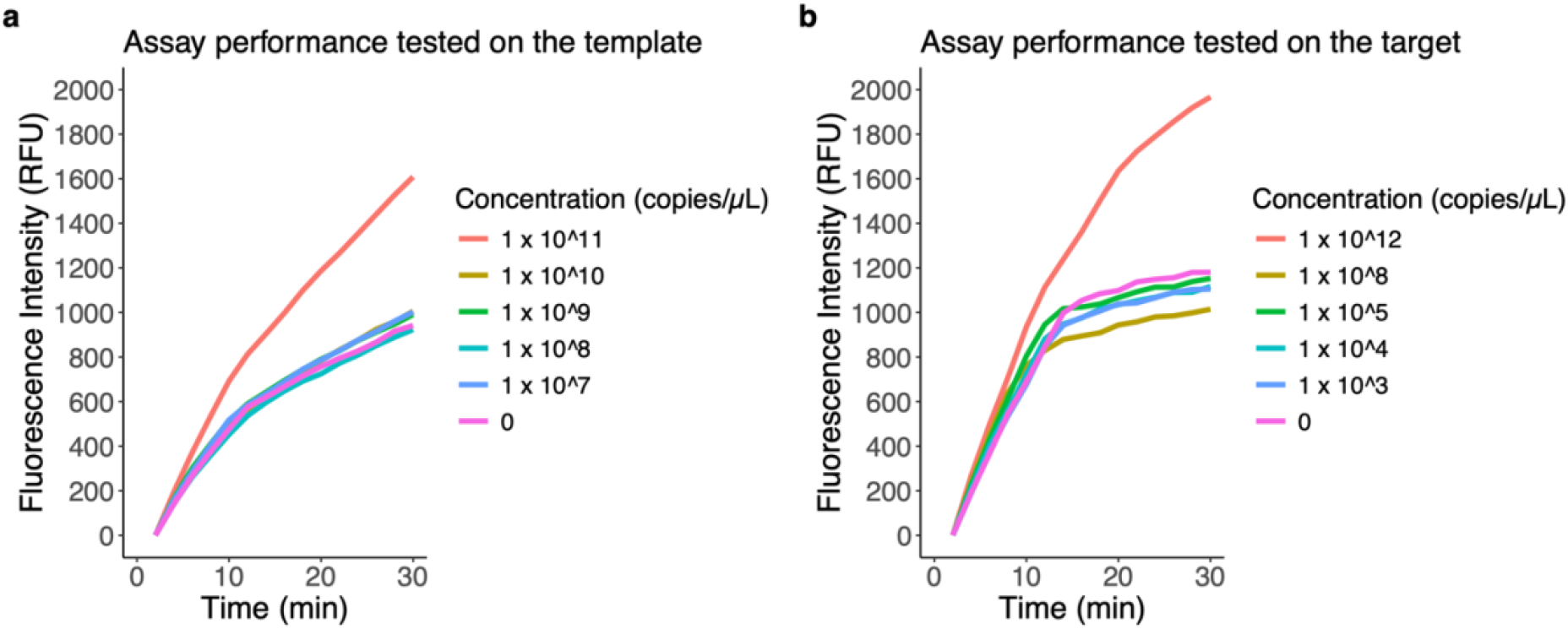
Fluorescence intensity when applying the NISDA assay on five template (a) or target (b) dilution series and a negative control (no template reaction). The assay was performed in TES buffer (pH 7.8) at a constant temperature of 42°C, with the fluorescence intensity of 6-FAM being recorded every 2 minutes. The published linear range for the assay on this template ranges from 10^-15 Molar to 10^-6 Molar; corresponding to 6.02 x 10^2 to 6.02 x 10^11 copies/µL. The average of two independent experiments is shown.

Previous studies on non-enzymatic nucleic acid detection methods have highlighted the importance of probe purity to prevent background leakage (noise signal), which may induce fluorescence even in the absence of the target^5–7^. While all probes were HPLC purified by the probe vendor, we assessed the purity of all reaction components using denaturing polyacrylamide gel electrophoresis (PAGE) (**Supplementary Fig. 3 and Supplementary Table 1)**. PAGE analysis verified the integrity of all reaction components but also revealed low abundant impurities, which were confirmed by high-performance liquid chromatography (HPLC) analysis (**Supplementary Fig. 4**). Therefore, an additional HPLC purification was performed to remove these impurities. We then repeated the NISDA assay on a template dilution series (as described in **Fig. 1a**), however, this did not improve results, and increased endpoint fluorescence was again only observed for template concentrations ≥ 1 x 10^11^ copies/µL.

In the original study, concerns were raised that hybridization of the recycled template with the M1:M2 duplex at lower reaction temperatures could impair signal amplification^4^. To address this, and to eliminate potential issues related to the temperature calibration of the Real-Time PCR instrument used in both the original study and our experiments, we conducted the assay at various temperatures using a template concentration of 1 x 10^11^ copies/µL. Additionally, we extended the reaction time to boost the amplification of the signal further. Despite these modifications, we were unable to discernably improve the difference in endpoint fluorescence between the template and no template reaction, as demonstrated in **Supplementary Fig. 5**.

The efficiency of this assay is primarily driven by free energy dynamics. The high background signal observed in our data suggests the possibility of a spontaneously initiated template-free reaction^8^. Therefore, we varied the concentration of the M1 and M2 probes to eliminate the possibility that inaccurate probe concentration measurements would impact reaction efficiency. Again, we did not observe an impact on endpoint fluorescence, as depicted in **Supplementary Fig. 6**. As the self-assembly of the M1 and M2 DNA probes is also driven by free energy, and the reaction efficiency could be limited by the spatial structures of these probes^9^, three distinct molecular beacon folding protocols (**Materials and methods**) were examined but these did not impact the reaction outcome (**Supplementary Fig. 7**).

Variations in reaction temperature and M1 and M2 probe concentration were also tested using a full factorial design (i.e. combining a temperature and probe concentration range). None of the tested conditions improved the outcome of the reaction (**Supplementary Fig. 8**).

## Discussion

Despite extensive efforts, we were unable to reproduce the sensitivity of the NISDA assay as reported by Mohammadniaei et al.^1^. A detectable fluorescence signal was only observed at target concentrations ≥ 1 × 10^11^ copies/µL, well above the originally claimed limit of 10 copies/µL.

We systematically tested a broad range of variables, including probe concentrations, probe folding protocols, temperatures, and reaction times. These experiments were replicated across different laboratories and with probes from multiple suppliers. None of these adjustments yielded improved assay performance. High background fluorescence persisted, potentially due to spontaneous, template-independent reactions, as previously noted in studies on entropy-driven and hybridization-based amplification systems^8,9^.

Enzyme-free isothermal amplification strategies hold great promise for robust, low-cost diagnostics, especially in resource-limited or point-of-care settings^9,10^. However, our findings underscore the importance of independent validation. While the concept behind NISDA is compelling, we could not replicate the reported sensitivity using the published protocol or variations thereoff, limiting its current utility for sensitive nucleic acid detection.

## Acknowledgements

This work was supported by the Interuniversity Special Research Fund (iBOF/23/030). We would like to thank the SOCan consortium for fostering a collaborative environment and engaging in fruitful discussions.

## Author contributions

T.VDS, S.A, and P.M identified the challenges described in the manuscript. The main results were jointly elaborated by T.VDS, S.A, S.B, A.P, A.C, E.C., and P.M. T.VDS, S.A, and P.M. drafted the manuscript. All authors contributed to the discussion and commented on the manuscript.

## References

1. Dirks, R. M. & Pierce, N. A. Triggered Amplification by Hybridization Chain Reaction. PNAS vol. 101 www.pnas.orgcgidoi10.1073pnas.0407024101 (2004).

2. Yin, P., Choi, H. M. T., Calvert, C. R. & Pierce, N. A. Programming biomolecular self-assembly pathways. Nature 451, 318–322 (2008).

3. Couny, F., Benabid, F., Roberts, P. J., Light, P. S. & Raymer, M. G. Generation and photonic guidance of multi-octave optical-frequency combs. Science (1979) 318, 1118–1121 (2007).

4. Mohammadniaei, M. et al. A non-enzymatic, isothermal strand displacement and amplification assay for rapid detection of SARS-CoV-2 RNA. Nat Commun 12, (2021).

5. Chen, X., Briggs, N., McLain, J. R. & Ellington, A. D. Stacking nonenzymatic circuits for high signal gain. Proc Natl Acad Sci U S A 110, 5386–5391 (2013).

6. Xiong, E., Yao, D., Ellington, A. D. & Bhadra, S. Minimizing Leakage in Stacked Strand Exchange Amplification Circuits. ACS Synth Biol 10, 1277–1283 (2021).

7. Sugawara, T. & Oishi, M. Latent Toehold-Mediated DNA Circuits Based on a Bulge-Loop Structure for Leakage Reduction and Its Application to Signal-Amplifying DNA Logic Gates. ACS Appl Mater Interfaces 16, 15907–15915 (2024).

8. Li, Y. et al. Entropy driven circuit as an emerging molecular tool for biological sensing: A review. TrAC - Trends in Analytical Chemistry vol. 134 (2021).

9. Chai, H., Cheng, W., Jin, D. & Miao, P. Recent Progress in DNA Hybridization Chain Reaction Strategies for Amplified Biosensing. ACS Applied Materials and Interfaces vol. 13 38931–38946 (2021).

10. Wang, W., Ge, Q. & Zhao, X. Enzyme-free isothermal amplification strategy for the detection of tumor-associated biomarkers: A review. TrAC - Trends in Analytical Chemistry vol. 160 (2023).

