## Supplementary information for "Non-enzymatic isothermal strand displacement and amplification (NISDA) does not enable sensitive nucleic acid quantification"

### Supplementary Fig. 1

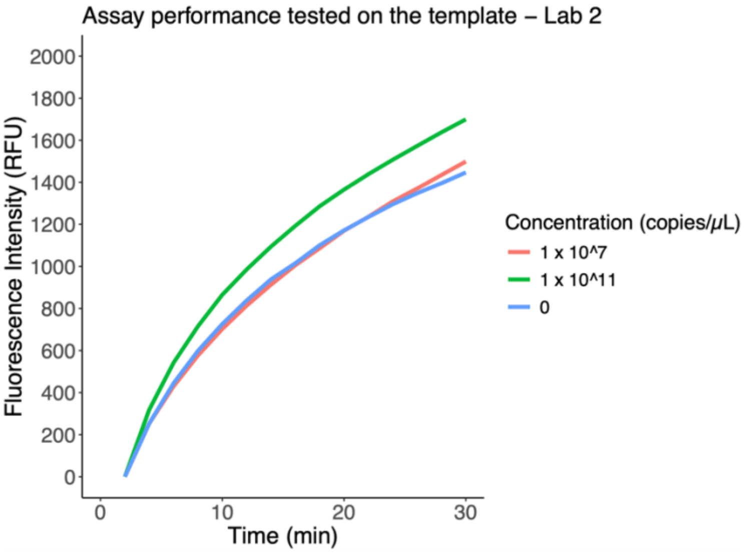


***Supplementary Fig. 1 Fluorescence intensity when applying the NISDA assay on two dilution points of the template and a negative control (no template reaction).*** *The assay was performed in TES buffer (pH 7.8) at a constant temperature of 42°C, with the fluorescence intensity of 6-FAM being recorded every 2 minutes. This experiment represents an independent repeat from the experiment shown in* ***Fig. 1*** *by another operator in another lab.*

### Supplementary Fig. 2

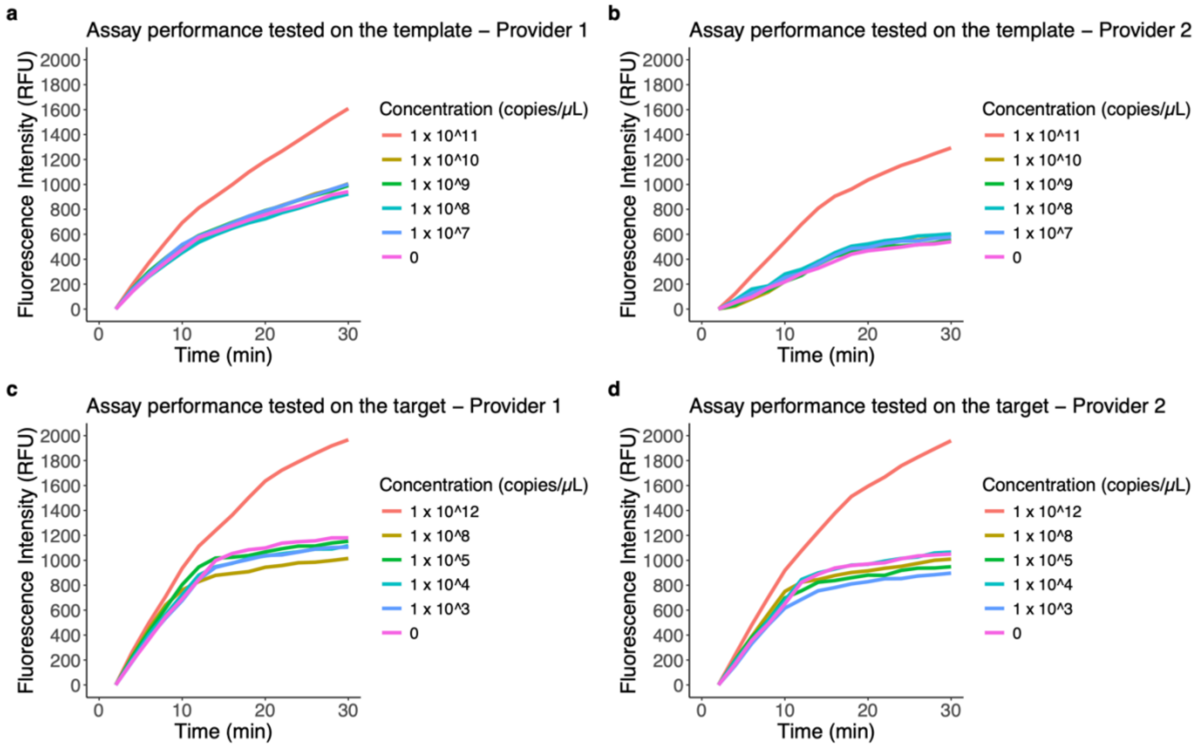


***Supplementary Fig. 2 Fluorescence intensity when applying the NISDA assay on five template (a,b) or target (c,d) dilution series and a negative control (no template reaction).*** *The assay was performed in TES buffer (pH 7.8) at a constant temperature of 42°C, with the fluorescence intensity of 6-FAM being recorded every 2 minutes. For* ***(a,b)*** *consumables from Integrated DNA Technologies (IDT) were used, and for* ***(c,d)*** *from the supplier mentioned in the original paper, Pentabase A/S. The average of two independent experiments is shown.*

### Supplementary Fig. 3

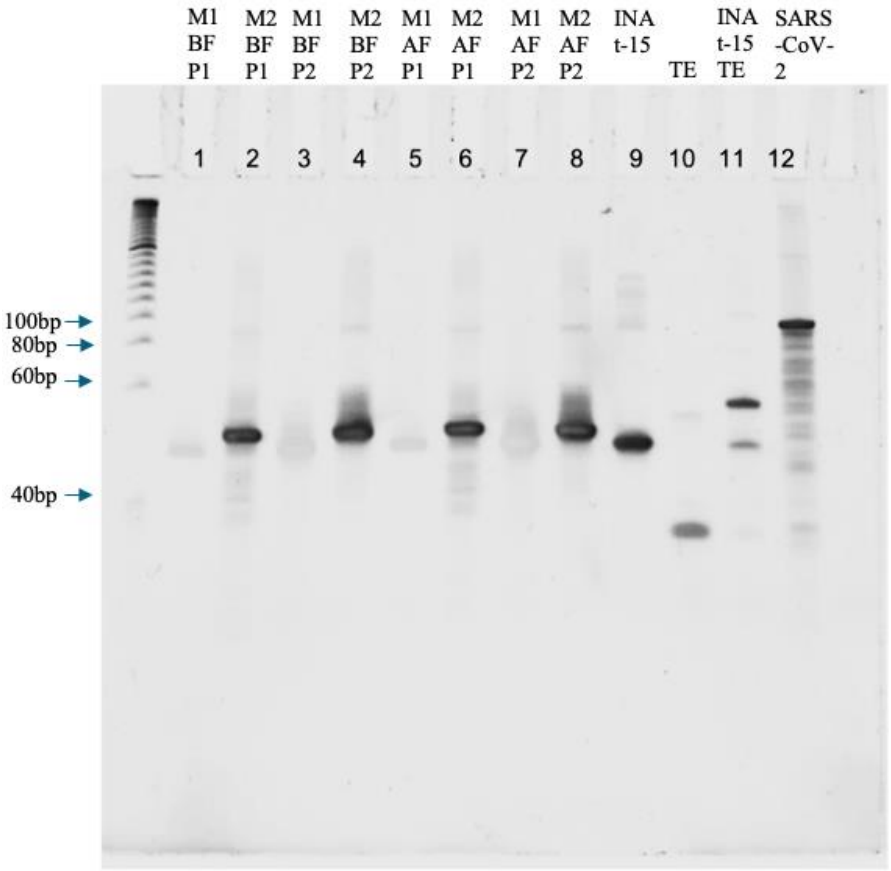


***Supplementary Fig. 3 A PAGE analysis with a gel containing: 12% acrylamide, 7 M Urea, and 1X Tris Borate EDTA buffer, pH 8.5****. The different lanes consist of: 1. the M1 probe before folding provided by IDT, 2. the M2 probe before folding provided by IDT, 3. the M1 probe before folding provided by Pentabase A/S, 4. the M2 probe before folding provided by Pentabase A/S, 5. the M1 probe after folding provided by IDT, 6. the M2 probe after folding provided by IDT, 7. the M1 probe after folding provided by Pentabase A/S, 8. the M2 probe after folding provided by Pentabase A/S, 9. the initiator sequence provided by Pentabase A/S, 10. The template sequence provided by IDT, 11. template/initiator complex after folding, and 12. the N-gene target sequence provided by IDT. M1 = the M1 probe, M2 = the M2 probe, BF = before the probe folding step, AF = after the probe folding step, INA t-15 = the 15nt long initiator sequence, including 4 intercalating nucleotides provided by Pentabase, TE = Template sequence, SARS-CoV-2 = the synthetic SARS-CoV-2 target mimic. The samples in lanes 1, 3, 5 and 7 have a lower response to the staining, compared to the other lanes.*

### Supplementary Table 1

***Supplementary Table 1:*** *The sequences of the oligonucleotides used.*

| Oligo Name | Sequence | Length |
|---|---|---|
| M1 probe | [FAM]_ATTGGCACCCGCAATAGGATTAATAGCAGGATTGCGGGTGCCAAT_[BHQ1] | 45nt |
| M2 probe | AGGATTAATAGCAGGATTGGCACCCGCAATCCTGCTATTAATCCTATTGCGGG | 53nt |
| INAt-15 | TAGCAGGATTGCGGGTGCCAATGTGATCTTTTGGTGT | 37nt |
| Template | ATTGGCACCCGCAATCCTGCTA | 22nt |
| Mimic-SARS-CoV-2 | GGTTGCAACTGAGGGAGCCTTGAATACACCAAAAGATCACATTGGCACCCGCAATCCTGCTAACAATGCTGCAATCGTGCTACAA | 85nt |

### Supplementary Fig. 4

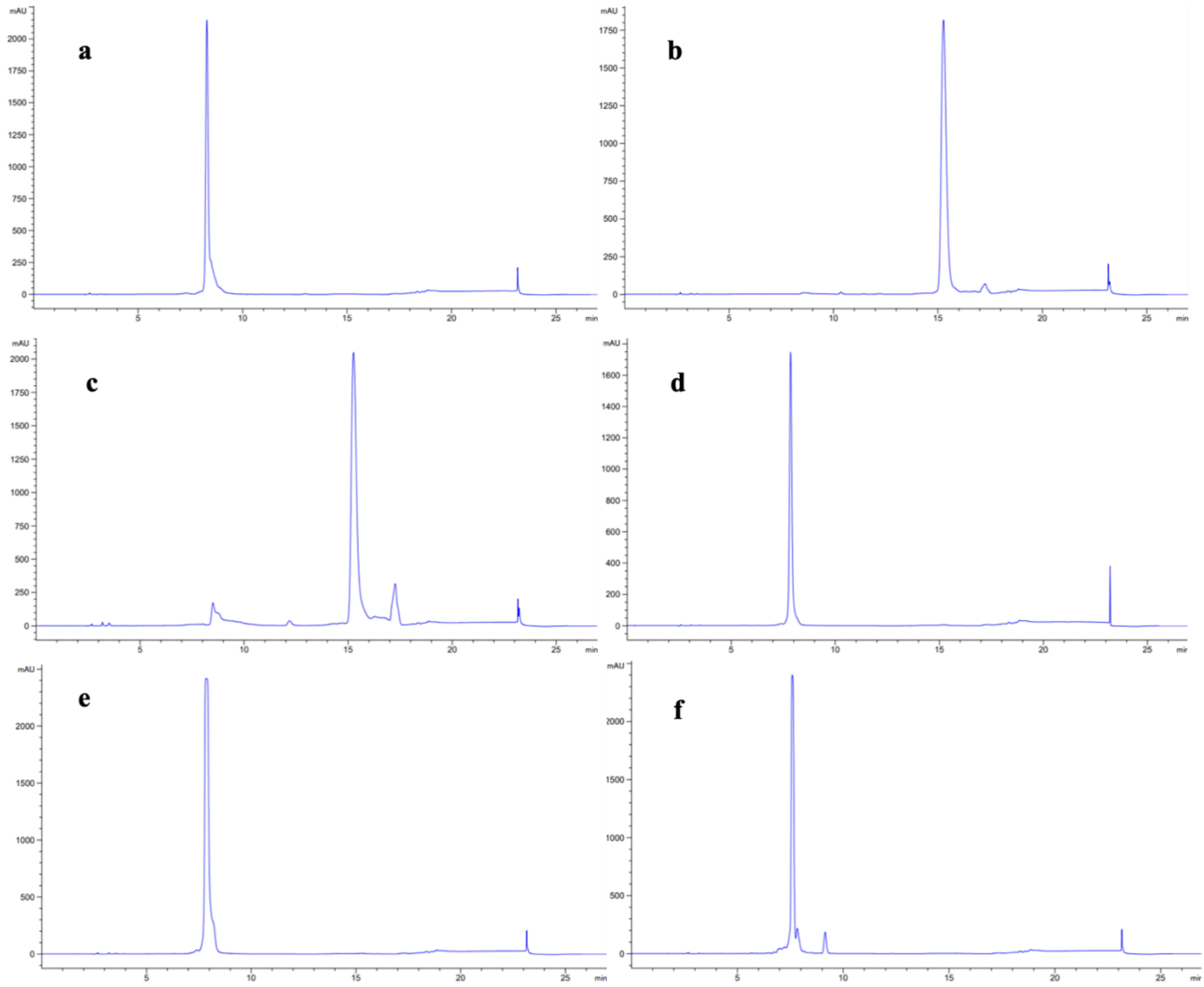


***Supplementary Fig. 4 The chromatograms after the HPLC analysis with the retention time (minutes) on the x-axis, and the detector response in milli-absorbance units (mAU) at a wavelength of 260nm on the y-axis. a.*** *the chromatogram of the initiator sequence.* ***b.*** *the chromatogram of the M1 probe provided by IDT.* ***c.*** *the chromatogram of the M1 probe provided by Pentabase A/S.* ***d.*** *the chromatogram of the M2 probe provided by IDT.* ***e.*** *the chromatogram of the M2 probe provided by Pentabase A/S, and* ***f.*** *the chromatogram of the template sequence.*

### Supplementary Fig. 5

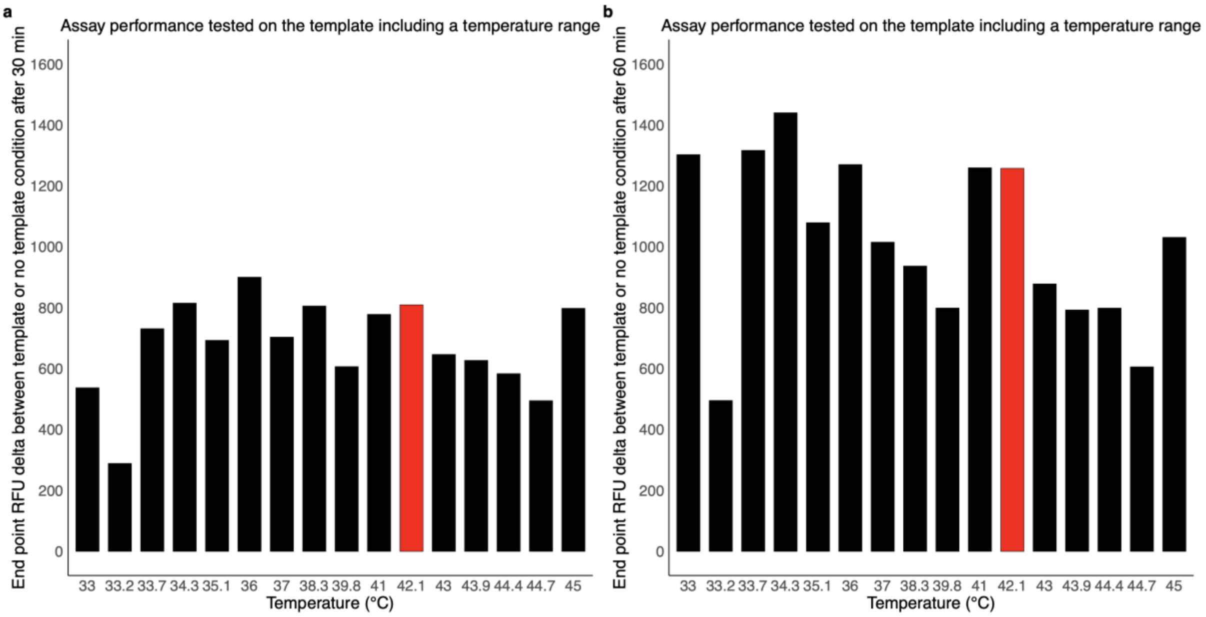


***Supplementary Fig. 5 Fluorescence intensity when applying the NISDA assay on one dilution point of the template (1 x 10^11 copies/µl) and a negative control (no template reaction).*** *The assay was performed in TES buffer (pH 7.8) at 16 different temperatures, ranging from 33°C until 45°C, with the fluorescence intensity of 6-FAM being recorded every 2 minutes for a duration of* ***a.*** *30 minutes and* ***b.*** *60 minutes. In the original paper a reaction temperature of 42°C is described as indicated by the red bars on the figures.*

### Supplementary Fig. 6

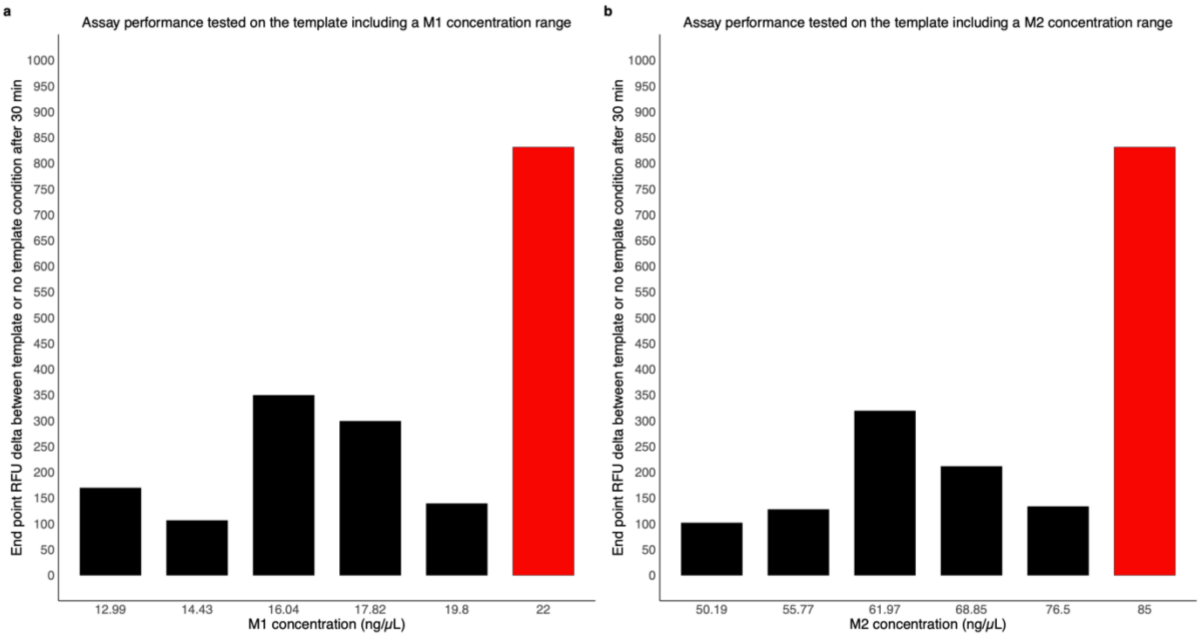


***Supplementary Fig. 6 Fluorescence intensity when applying the NISDA assay on one dilution point of the template (1 x 10^11 copies/µl) and a negative control (no template reaction).*** *The assay was performed in TES buffer (pH 7.8) at a constant temperature of 42°C, with the fluorescence intensity of 6-FAM being recorded every 2 minutes for a duration of 30 minutes. In* ***a.*** *the concentration of M1 was varied and in* ***b.*** *the concentration of M2 was varied. In the original paper a M1 concentration of 22 ng/µL and M2 concentration of 85 ng/µL is described as indicated by the red bars on the figures.*

### Supplementary Fig. 7

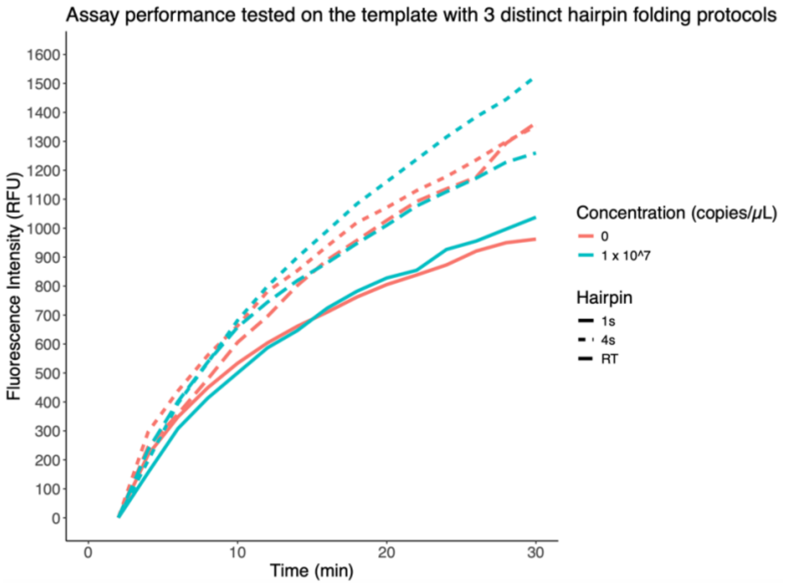


***Supplementary Fig. 7 Fluorescence intensity when applying the NISDA assay on one dilution point of the template (1 x 10^11 copies/µl) and a negative control (no template reaction).*** *M1 and M2 probes were folded using three different methods: 1) RT: incubation at 95°C for 5 minutes followed by cooling to room temperature, 2) 1s: incubation at 95°C followed by ramped cooling to room temperature at 0.1°C/s and a 1-hour incubation at room temperature, and 3) 4s: incubation at 95°C followed by ramped cooling to room temperature at 0.1°C/s with intermediate stops at 77°C, 59°C, and 41°C for 12 minutes each. In the original paper, the following protocol is described: the probes M1 and M2 in TES buffer (pH 7.8) are formed by annealing at 95°C for 5 min followed by a gradual cooling down to room temperature (RT) for 1 h.*

### Supplementary Fig. 8

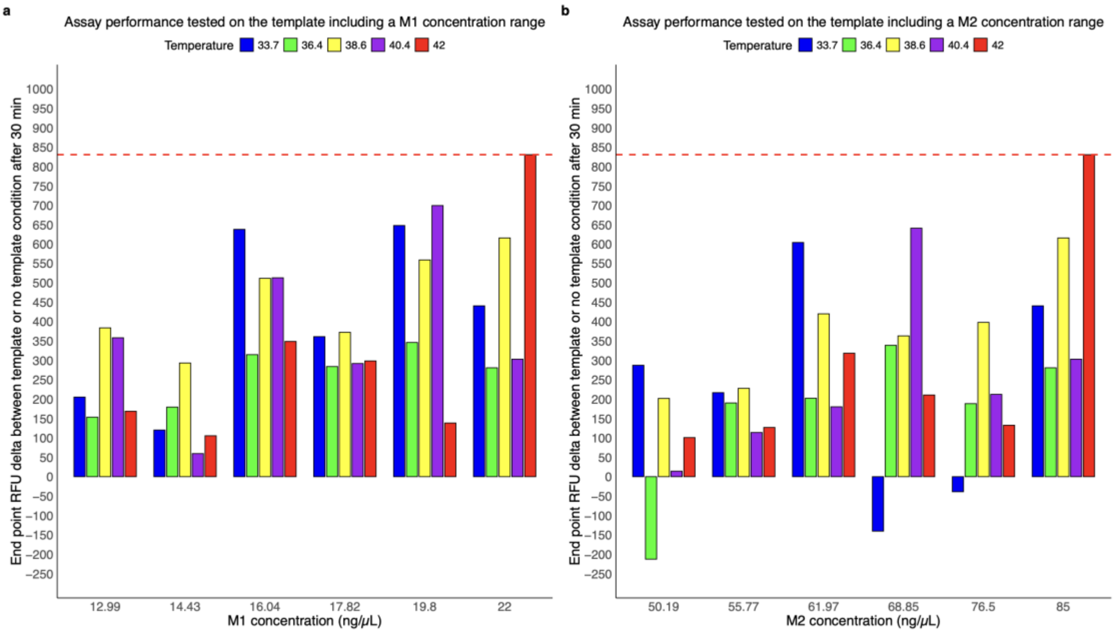


***Supplementary Fig. 8 Fluorescence intensity when applying the NISDA assay on one dilution point of the template (1 x 10^11 copies/µl) and a negative control (no template reaction).*** *The assay was performed in TES buffer (pH 7.8) at five different temperature points (33.7°C, 36.4°C, 38.6°C, 40.4°C, and 42°C), with the fluorescence intensity of 6-FAM being recorded every 2 minutes for a duration of 30 minutes. In* ***a.*** *the concentration of M1 was varied and in* ***b.*** *the concentration of M2 was varied. In the original paper a M1 concentration of 22 ng/µL, a M2 concentration of 85 ng/µL, and 42°C as incubation temperature is described as indicated by the red dashed lines on the figures.*
